# Comparative transcriptomic response of bank vole kidney and respiratory epithelial cells to distinct orthohantaviruses

**DOI:** 10.64898/2026.11.11.765681

**Authors:** Karine Badonnel, Thomas Cokelaer, Elodie Turc, Marc Monot, Jean Claude Manuguerra, Myriam Ermonval, Giulia Gallo

## Abstract

Orthohantaviruses are rodent-borne viruses distributed worldwide that can cause diseases of varying severity in humans following occasional aerosol transmission. While infections remain asymptomatic in their natural small-mammal reservoirs, hantaviral pathogenesis in humans is associated with an exacerbated immune response. The increasing number of outbreaks in endemic regions caused by the modification of rodent habitats due to human activities and the risks of reassortments in areas where multiple orthohantaviruses co-circulate with their animal reservoirs, are of public health concern. These challenges highlight the need to better understand how these viruses interact with host cells to spread within and between organisms, and how antiviral responses are regulated, particularly in their natural hosts. We previously showed that cellular susceptibility to orthohantaviruses depends on both cell type and virus species and correlates with cellular restrictions acting at different stages of the viral cycle. Here, we performed a comparative RNA-sequencing study and transcriptomic analysis of the cellular responses induced by three different orthohantaviruses in kidneys and respiratory tract cells derived from bank voles, the natural host of an orthohantavirus pathogenic to humans. Transcriptomic analyses revealed common as well as virus– and tissue-specific regulatory pathways that may underlie distinct infection outcomes. It highlighted the complexity of viral interactions with bank vole at a cellular level, including innate immune responses and antiviral signaling pathways.

## Introduction

Orthohantaviruses (1) are found in small mammals, such as rodents, shrews, moles and bats. Their long co-evolution with these globally distributed host species (2–4) accounts for their presence in all continents (5). These viruses do not cause apparent diseases in their natural hosts. Currently, only rodent-borne orthohantaviruses are known to be pathogenic in humans, mainly through the inhalation of aerosols from contaminated animal excreta. After entering the respiratory tract, orthohantaviruses cross the pulmonary epithelial barrier and disseminate throughout the body by infecting vascular endothelial cells and monocytes (6, 7). In humans, an exacerbated immune response (8) leads to hemorrhagic fevers with renal syndromes (HFRS) and nephropathia epidemica (NE) in the Old World (9), or to hantavirus cardio-pulmonary syndromes (HCPS) in the Americas (10). The activation of innate immunity to eliminate viruses, and the viral strategies to counteract it, vary depending on the orthohantavirus species (11). Sustained interferon (IFN) activation may contribute to inflammatory responses that influence viral pathogenicity (12).

We previously showed that the pathogenic Puumala virus (PUUV), the weakly pathogenic Tula virus (TULV) and the non-pathogenic Prospect Hill virus (PHV), differentially regulate IFN responses in human and rodent cells (13, 14). *Myodes glareolus* (bank vole) is the natural host of PUUV, whereas TULV and PHV naturally infect *Microtus arvalis* (common vole) and *Microtus pensylvanicus* (eastern meadow vole), respectively. These orthohantaviruses exhibit different replication characteristics in bank voles-derived cells. These differences correlate with cellular restrictions acting at distinct stages of the viral cycle from entry to replication and viral release (14). In human cells, type I and III IFNs and several IFN stimulated genes (ISGs) are strongly activated by PUUV and PHV but not by TULV. In contrast, bank vole cells display a delayed and weaker IFN response.

Despite advances in our understanding of the regulation of host responses in rodents, the mechanisms leading to the differential outcomes of orthohantavirus infection in their natural hosts remain poorly understood (15–18). To explore this question, we compared gene expression profiles induced by PUUV, TULV and PHV. Building on the opportunity that the recent availability of an updated *Myodes glareolus* genome assembly in the NCBI database (<u>GCF_902806735.1</u>) provides, we performed transcriptomic analysis by RNA sequencing (RNA-seq) of bank vole epithelial cells derived from the kidneys (MyglaSWRecB) and the respiratory tract (MyglaSWTrach and MyglaAEC#2) infected with these different orthohantaviruses. Differential gene expression and pathway analyses revealed virus– and tissue-specific regulatory programs that likely contribute to distinct infection outcomes and host–virus interactions.

## Material and methods

### Cells and viruses

The three bank vole cells lines, MyglaSWRecB, MyglaSWTrach, MyglaAEC, originated from kidney, tracheal and alveolar lung epithelium, respectively, and were SV40 T antigen-immortalized (19). Cells were grown at 37°C with 5% CO2 in DMEM medium (Gibco) supplemented with 10% of heat-inactivated fetal bovine serum (Biosera).

Puumala Sotkamo viral strain (PUUV) and Prospect Hill viral strain (PHV) were provided by Andreas Rang (Charité University, Berlin). Tula Moravia virus (TULV) was obtained from Alexander Plyusnin (Helsinki University). Viral stocks were propagated on Vero E6 cells as previously described (14).

### RNA preparation, sequencing and transcriptomic analysis

RNA preparation from infected cells was performed as previously described (14). Briefly, bank vole cells were infected with orthohantaviruses at a multiplicity of infection (MOI) of 0.5. Total RNA was extracted 5 days post infection (dpi). After washing with PBS, cells were treated with Trizol (Tri Reagent, Sigma Aldrich) and chloroform. The aqueous phase containing the RNA was precipitated with isopropanol and resuspended in distilled water. RNA concentration was measured using a NanoDrop spectrophotometer (Thermo Scientific). Four biological replicates were generated for each condition: non-infected controls (NI) and cells infected with PUUV, PHV or TULV orthohantaviruses.

Strand-specific single-end cDNA libraries were prepared from total RNA using the Illumina Stranded mRNA prep, ligation preparation kit (Illumina) according to the manufacturer’s instructions. Libraries were sequenced on an Illumina NextSeq 2000 platform.

RNA-seq analyses were performed using the Sequana framework, v0.14.1 (20). Viral infection in samples was first verified using the Sequana mapper workflow to detect specific viral RNA sequences. Transcriptomic analysis was conducted using the Sequana RNA-seq pipeline (v0.15.2), implemented in Snakemake (21) and available at https://github.com/sequana/sequana_RNA-seq, using default parameters. Adapter trimming and quality filtering were performed with fastp v0.20.1 (22). Cleaned reads were aligned with STAR (23) against the *Myodes glareolus* reference genome assembly GCA_902806735.1 and the *Mus musculus* reference genome (GRCm38/mm10). Library strandedness was automatically inferred by the pipeline, and gene-level counts were generated using FeatureCounts (package Subread v2.0.1). Differential gene expression analysis was performed from the count matrix using DESeq2 (v1.38.2) with R v4.2.1. Statistical significance was assessed using Benjamini–Hochberg adjusted p-values with a false discovery rate (FDR) threshold of 0.05.

All third-party software dependencies were executed within containerized environments provided through the Damona project (https://damona.readthedocs.io), ensuring reproducibility.

### Visualization of differentially expressed genes

Venn diagrams were drawn using the online Venny 2.1 program, an interactive tool for comparing lists with Venn’s diagrams (24) allowing the determination of the number and identification of genes common or specific to a given situation. Volcano plots and gene comparison representations were obtained using GraphPad Prism 11. Heatmaps representations of differentially expressed genes (DEGs) were generated using the pheatmap function in the R program. This analysis was performed for tracheal cells infected with PUUV, PHV, or TULV across the bank vole and mouse reference genomes (BV and mm10). For each condition, only one entry per gene was retained, prioritizing the lowest adjusted p-value (padj) and, in case of ties, the largest absolute log2FoldChange. For the relative representation of genes identified through the BV genome in comparison to the mm10 genome specifically in infected tracheal cells, the six DEGs tables were merged by gene union, with missing values replaced by zero. Genes with non-finite values (NA, NaN, Inf) or zero variance across conditions were removed. Up and downregulated genes were defined based on the sign of the mean log2FoldChange across the six conditions. Heatmaps were generated using pheatmap in R, scaling values by gene (scale = “row”), clustering columns using Euclidean distance and complete linkage, and annotating columns by virus and genome. This approach enabled visualization of global expression profiles and comparison of regulatory patterns across viruses and genomes.

### Pathway enrichment

Functional enrichment analysis of differentially expressed genes (DEGs) was performed using DAVID Bioinformatics Resources v6.8 (https://davidbioinformatics.nih.gov/tools.jsp) (Database for Annotation, Visualization and Integrated Discovery (25). Official gene symbols were submitted for annotation clustering and enrichment analyses of Gene Ontology (GO) terms, KEGG and Reactome pathways, and InterPro protein domains. The background dataset was set to the entire *Mus musculus* genome, and Enriched terms were considered statistically significant based on a Benjamini-Hochberg adjusted p-value < 0.05.

## Data availability

The RNA-seq data set 1 was published in (14) with accession GSE198751. The RNA-seq data set 2 and 3 were deposited to ENA (accession E-MTAB-16656).

## Results

### Quantitative gene regulation in kidney cells or in respiratory cells varies depending on orthohantavirus species

To evaluate the effect of orthohantavirus infection on gene expression, different epithelial bank vole (BV) cell lines derived from organs targeted by these viruses, such as the respiratory tract (MyglaSWTrach and MyglaAEC#2) and the kidneys (MyglaSWRecB), were infected at a MOI of 0.5 with PUUV, TULV or PHV and their RNAs extracted five days post infection (5 dpi). At this time point, the viral replication of PUUV and PHV is clearly detectable and constitutes a suitable basis for the quantitative and qualitative characterization of gene expression by RNA-seq, as previously described (14). The experimental approach is schematized in Figure 1. The RNA-seq generated a similar number of reads (20-30 million) to each sample and these samples are clustered according to infection status distinguishing uninfected and infected conditions (Fig. SD1). The number of identified genes significantly upregulated and downregulated (FC>2 and <-2, p-value <0.05), varied according to the virus and cell types (Table 1). Venn diagrams illustrate, in addition to the number of genes, the percentage of genes specific or common to different situations. Since few genes were upregulated upon orthohantavirus infection of pulmonary alveolar MyglaAEC#2 cells (Table 1), they were not included in Venn diagrams and volcano plot representations. Around 70% of differentially expressed genes (DEGs), were upregulated and 30% were downregulated. However, more genes appeared differentially expressed (DE) in the infected tracheal cells than in the kidney cells, both with PUUV (308 vs 66, Fig. 2A) and PHV (545 vs 254, Fig. 2B). Strikingly, few DEGs were common to these two cell lines upon PUUV infection (1.1%) or PHV infection (8.1%). In the case of TULV (Table 1) very few genes were DE in MyglaSWRecB compared to what is seen in MyglaSWTrach (4 vs 509).

**Figure 1:**
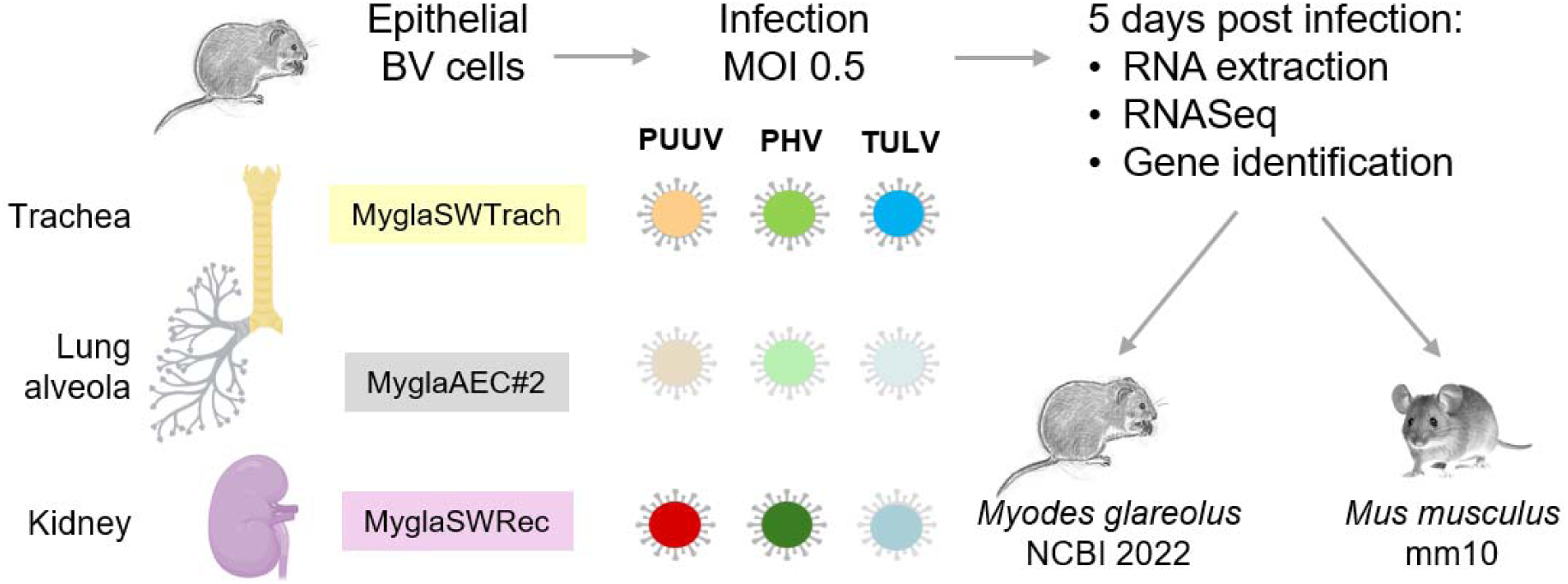
Schematic representation of the experimental protocol. As described in material and method, epithelial cells originated from different bank vole organs. Cells derived from the respiratory tract, MyglaSWTrach from trachea and MyglaAEC#2 from lung alveola, or from kidneys, MyglaSWRecB, were infected with Puumala, Prospect Hill and Tula orthohantaviruses (PUUV, PHV, TULV, respectively). Four samples were prepared per condition. RNA was extracted 5 days post infection and sequenced for gene identification on both bank vole (BV-NCBI 2022) and mouse (mm10) genome databases. Organs were drawn using BioRender.

**Figure 2:**
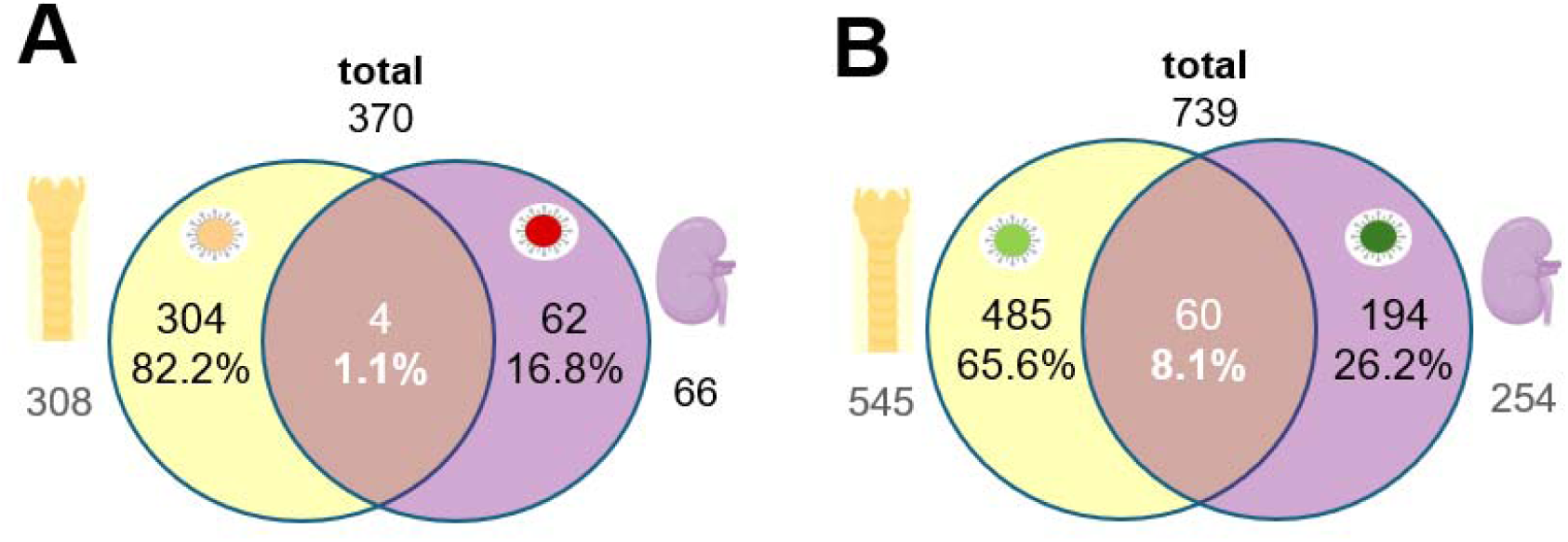
Regulation of genes in different bank vole cells. The Venn diagrams show the number of genes significantly regulated by PUUV (A) and PHV (B) in the different BV cell lines MyglaSWTrach (in yellow), or MyglaSWRec (in violet).

While TULV did not induce significant gene regulation in the kidney cell line, PUUV and PHV did (Fig. 3A). Indeed, in MyglaSWRecB cells, 37 of 66 genes (56%) were specifically regulated by PUUV, and 226 of 254 genes (88.9.%) by PHV. This represented 12.5% and 76.6% of the total identified genes for PUUV and PHV respectively, when only 9.5% of genes were common to PUUV– and PHV-infected cells. In the tracheal cells, MyglaSWTrach, the situation was quite different (Fig 3B). Many more genes were DE upon infection with the nonpathogenic orthohantaviruses, PHV (545) and TULV (509), than with PUUV (308). Among these genes, only 10.5% were specific to PUUV infection, 21.6% to PHV infection and 15.4% to TULV infection. Interestingly, in this cell line, a higher proportion of DEGs was shared following infection with all three orthohantaviruses (21.2%) or with the non-pathogenic heterologous orthohantaviruses PHV and TULV (23.7%). In contrast, fewer genes were common to PUUV and PHV or PUUV and TULV (3.1% and 4;6% respectively). Although the volcano plots show gene expression levels (log2FoldChange) and their significance (-log10_adjusted p-value), it should be noted that more genes were upregulated than downregulated in each situation (Fig. 3C).

**Figure 3:**
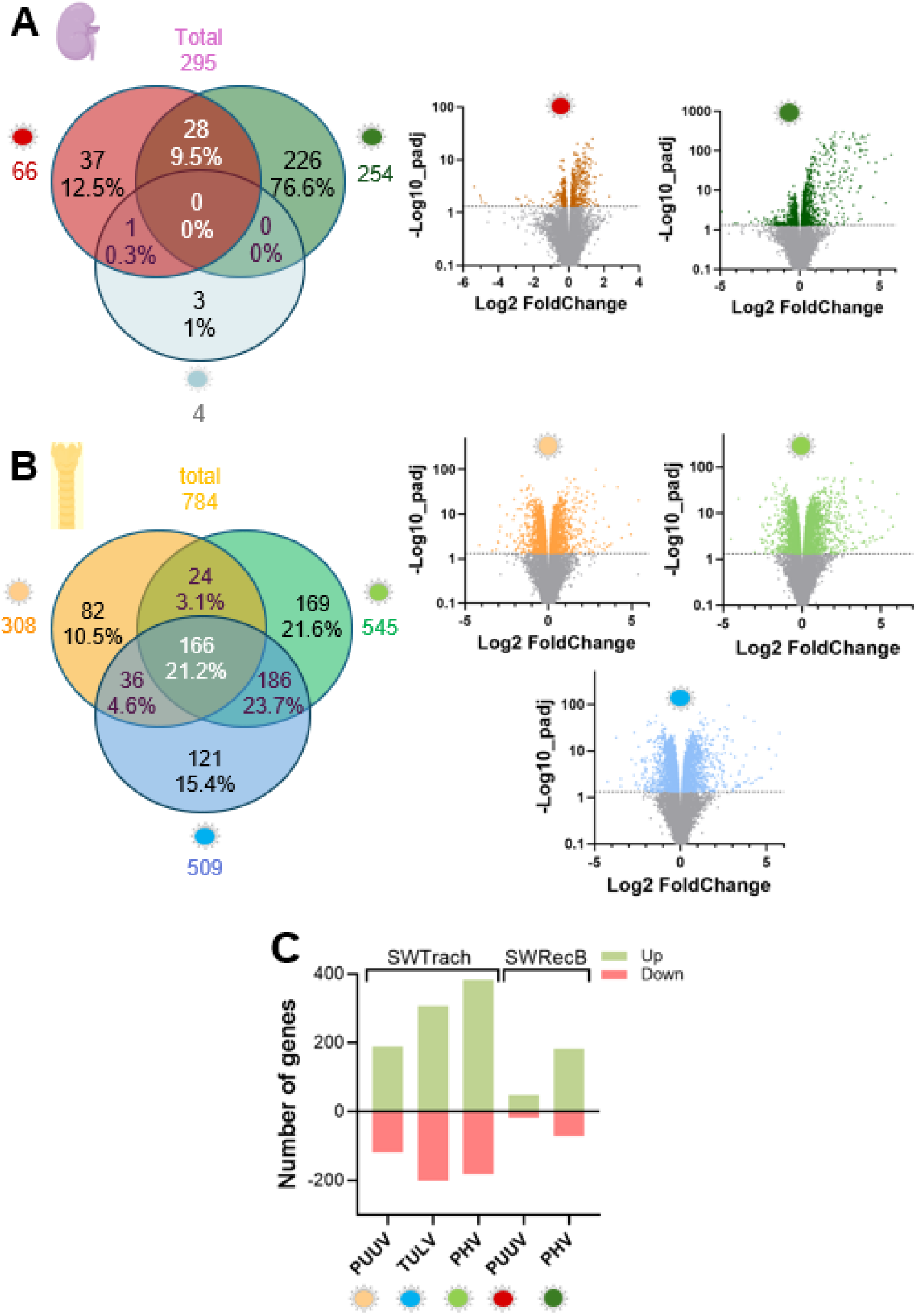
Regulation of genes by different orthohantaviruses. The Venn diagrams (left panels) show the number of DEGs significantly regulated in the different BV cell lines, MyglaSWRecB (A) and MyglaSWTrach (B) infected either with PUUV (orange), PHV (green) or TULV (blue). The distribution (x axis) and significance (y axis) of up and down regulated genes by the different orthohantaviruses are shown in the volcano plots (right panels) for MyglaSWRecB (A) and MyglaSWTrach (B) respectively. The histogram (C) summarizes the number of up and downregulated genes in the different conditions of infection.

Due to the limited information available for the bank vole genome, our previous work identified DEGs in infected renal cells from the mouse database (mm10). Therefore, the RNA-seq data generated in the present study were also analyzed against this mouse database. The percentage of DEGs found in infected tracheal (Fig. SD2A) or renal (Fig. SD3A) cells, specific or common to different viruses was similar, whether using the bank vole or the mouse reference genome. However, it is worth noting that for PUUV, PHV and TULV, only about 20% of the identified genes in MyglaSWTrach cells were found with the two genomes (Fig. SD2B), highlighting the importance of combining the two analyses. The same observation was made for the renal cells (Fig. SD3B), where an even smaller percentage of genes appeared common to the two databases. This result should nevertheless be viewed with caution due to the small number of DEGs identified after PUUV infection of MyglaSWRecB cells. Therefore, all the annotated genes found in the mouse and vole reference genomes were compiled into a single list of identification (Table 2 and Table SD). It should also be noted that many genes annotated as genomic loci (Loc genes) have orthologues in the mouse or human genome. This is for instance the case for 25, 54 and 42 Loc genes (FC>4 and <-4), for PUUV, PHV and TULV respectively in tracheal infected cells, and for 7 and 72 genes, for PUUV and PHV in renal infected cells. The putative corresponding names of these identified sequences have been added to the tables (in blue) and when several sequences referred to a similar orthologue, only the one with the highest FC value was retained.

### Innate immune responses are activated by orthohantaviruses in kidney and respiratory epithelial cells from bank vole

We compared the DEGs identified in the renal and the tracheal cell lines infected with the different orthohantaviruses, PUUV, PHV, TULV, using the bank vole and the mouse reference genomes (Table 2). The transcripts regulated by one or more viruses in different BV cell lines (Fig. 4A) were mainly involved in antiviral defense as expected following the recognition of RNA viruses by Toll-, Nod– or Rig1-like receptors (TLR, NLR, RLR) upstream of innate immune responses (12). Only two genes were significantly activated by PUUV (FC>2) in both MyglaSWTrach and MyglaSWRecB cell lines; Ddx60, a helicase promoting RLR signaling, and the IFN-inducible gene Ifit1, known to inhibit viral replication (26). These two genes were also upregulated by PHV in these two BV cell lines. It is remarkable that only a few ISGs were slightly upregulated by PUUV, the orthohantavirus specifically transmitted by bank voles, as compared with the much stronger induction observed following infection with the heterologous orthohantavirus PHV. Additional genes upregulated by PHV only in the renal and the tracheal cell lines are shown in Figure 4B, and transcriptional variations induced by PHV in the two cell lines are represented in Figure 4C. Among the DEGs showing the highest fold changes and common to the two cell lines, we found genes involved in IFN signaling such as Oas1, Irf7, Zbp1, Oasl1, and several interferon-induced (IFI) genes or interferon-stimulated genes (ISGs) known as host restriction factors, including Rsad2, Bst2, Mx1, Mx2, Oas2, Ifi203, Ifi44L, as well as Usp18, a negative regulator of the IFN signaling. Other IFN-induced GTPase Tgpt2, Iigp1, Irgm1, Gpb4-6, the chaperone protein Rtp4 and the phospholipase Pla2g5 together with, at lower fold change levels, the proinflammatory cytokine Cxcl10, genes involved in nucleotide biosynthesis Cmpk2, immune regulation Lgals9, and the complement cascade Cfb and C2, were also activated. Fourteen of these genes (Table 2) were also slightly, but significantly (p-adjusted p-value <0.05), activated by PHV in the alveolar epithelial cells from lung, MyglaAEC#2. Some of the ISGs identified in tracheal and renal cells may have inhibitory effects on viral transcription and replication (e.g., Ifit, Mx1 and Cmpk2). Other DEGs could restrict viral entry, either at the level of receptor expression or fusion of the viral envelope with endo-lysosomal membrane (e.g., Samd9 and Ly6e). Others may affect virus release at the stage of vesicular budding (Rsad2/viperin (27)) promoting intracellular retention of viral particles (Bst2/tetherin (28)) or inducing autophagy (Zbp1), while the host restriction factors Cmpk2 and Rsad2 are thought to act synergistically (29). Of note, another pro-apoptotic gene, Xaf1 (30), with the highest fold change (FC >200) in A549 human cells infected with PUUV (14), was only slightly activated (FC=2) in renal cells infected with PHV. The galectin Lgals9, which binds terminal galactose on glycans and has multiple biological functions (31) could also contribute to inflammation and immune response during orthohantavirus infection.

**Figure 4:**
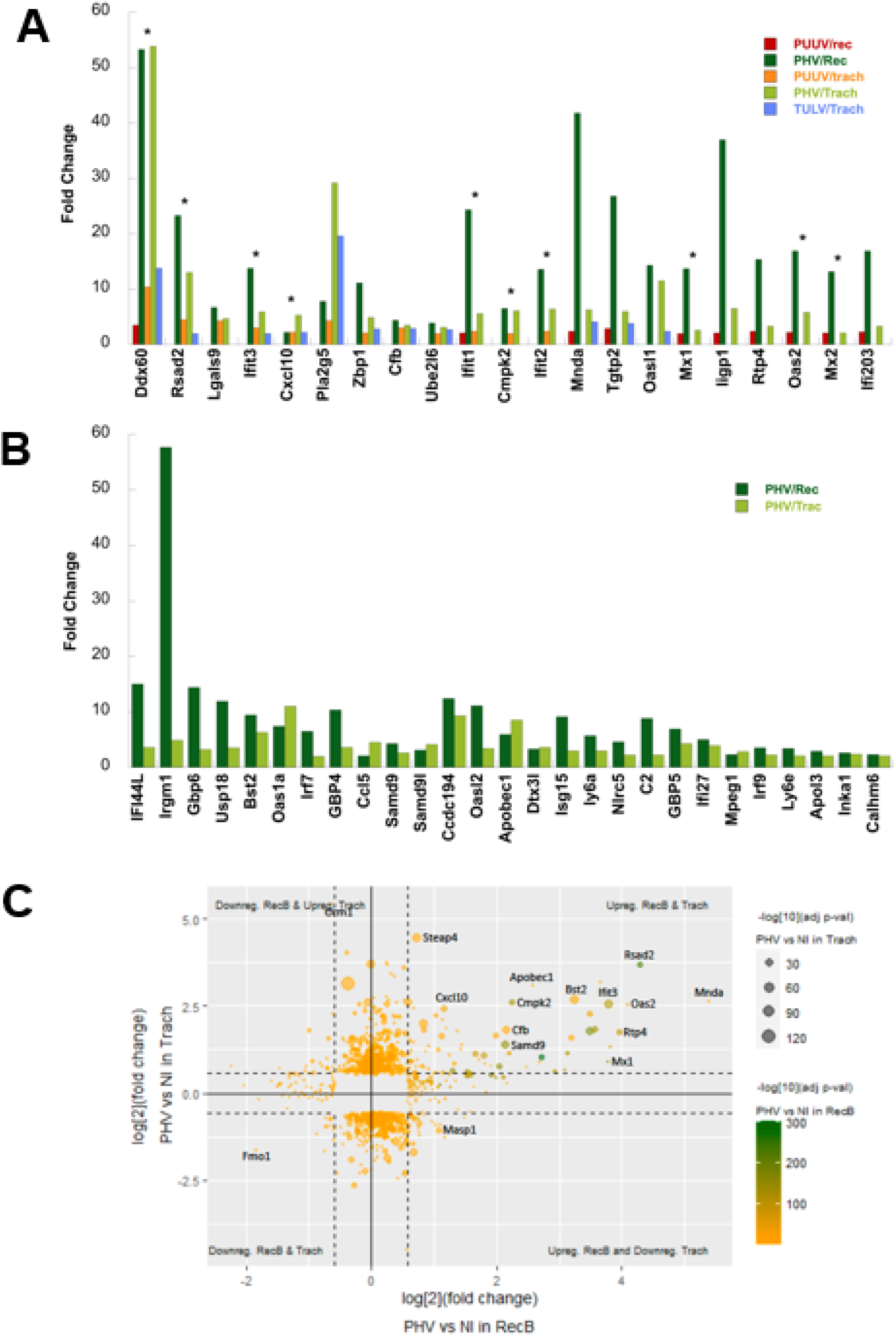
Genes common to different infected cell lines. The histogram (A) represents the variation of the fold change of genes significantly upregulated by PUUV and PHV in different cell lines as indicated in the color legend, and the asterisks show the genes previously found upregulated by PUUV in human epithelial lung cells (see table 2) The histogram (B) shows the fold change of the genes upregulated by PHV, both in renal (dark green bars) and in tracheal (light green bars) cells. The scatter plot (C) compares PHV-induced transcriptional responses between renal MyglaSWRecB (RecB) and tracheal MyglaSWTrach (Trach) cells. Only genes displaying significant differential expression in at least one cell type (|fold change| ≥ 1.5 and adjusted p-value < 0.05) were retained. It shows the log₂ fold change of gene expression in PHV-infected RecB cells versus non infected (NI) cells on the x axis and PHV-infected Trach cells versus NI on the y axis. Each point corresponds to a gene shared between the two datasets. Point color represents the statistical significance of differential expression in renal cells, whereas point size reflects the significance in tracheal cells. Quadrants distinguish genes showing similar regulation (matches) or opposite regulation (mismatches) between the two cell types. Solid lines mark zero fold change and dashed lines indicate the fold-change threshold (|FC| = 1.5).

Other genes involved in anti-viral defense were differentially expressed only in infected MyglaSWTrach cells. Among DEGs shared by the 3 viruses, PUUV, PHV, TULV (Table SD1A), or by 2 of them (Table SD1B), many were IFN-induced. This included proinflammatory cytokines and GTPase. It is noteworthy that the cytosolic IFN sensor Sting1, which was not differentially expressed in the other infected bank vole cell lines, was activated by the 3 orthohantaviruses in tracheal cells. This gene has been shown to regulate different aspects of cell death such as apoptosis and autophagy (32). An antiviral role of this pathway during Hantaan orthohantavirus (HTNV) infection via apoptosis was recently reported (33). Therefore, the STING pathway together with other regulators of apoptosis could be involved in orthohantavirus elimination by tracheal cells.

In addition, the innate immune response in MyglaSWTrach cells was characterized by specific upregulation of Ch25h, an IFN-induced gene, in cells infected with PHV and TULV. Ch25h is involved in cholesterol metabolism and has been shown to inhibit membrane fusion and therefore entry of some enveloped viruses (34, 35). Ifitm3, also known to inhibit viral entry (36) was slightly upregulated only in PHV-infected kidney cells. Different members of the tripartite motif-containing protein family (TRIM), which appear to be important players in viral defense (37) were DE either in tracheal or renal BV cells. Although exhibiting low fold changes, their expression varied depending on the orthohantavirus species. In tracheal cells, Trim15 was activated and Trim66 was downregulated by all three viruses while Trim10 was upregulated only by PUUV and PHV, and Trim9 was downregulated by PHV (Table SD1). Other members, Trim30, Trim34, Trim69, Trim14 and Trim21 were upregulated by PHV in renal cells (Table SD2), highlighting the complexity of TRIM family regulation. Of note, HTNV has been shown to inhibit the innate IFN response by targeting Trim25 (38).

In kidney cells, most of the PHV-induced antiviral ISGs originated from type I and III IFN responses (Table SD2). However, some guanylate-binding proteins (GBP), IFN-γ induced GTPases capable of interfering with multiple stages of the viral life cycles (39, 40), were specifically activated in renal cells. This included Gbp7, Vlig1, Gbp11, Gbp8 and Gbp1, which were not identified in tracheal cells, but have recently been described as restricting HTNV entry in human cells (41, 42)

Taken together, these results support that PHV, which is not specific to the bank vole, induced a stronger innate antiviral response than PUUV and that even in the context of anti-viral defense, different genes can be activated depending on the virus and the cell type.

### RNA-seq analysis and functional enrichment of DEGs in BV cell lines reveal cell type specificities of gene regulation by orthohantavirus infection

As showed above, more genes were upregulated and downregulated in the infected tracheal cells than in the infected kidney cells, and many of the identified genes in tracheal cells were not found in the kidney cells and vice versa (Table SD1 and SD2A). In contrast to genes of the antiviral response for which the FC values in PUUV-infected cells were often lower than those observed in PHV-infected cells (Table 2 and Fig 4), the DEGs specific to tracheal cells, either common to infections with all three orthohantaviruses, PUUV, PHV and TULV (Table SD1A), or shared by two of them (Table SD1B), exhibited similar FC ranges. It should be noted that in addition to genes regulated by the three viruses, only a few were shared by PUUV and PHV (26 genes) or PUUV and TULV (34 genes), whereas a much larger number of genes (179 genes) was common to the heterologous PHV and TULV orthohantaviruses. This observation is supported by the clustering of DEGs from PHV– and TULV-infected MyglaSWTrach cells, regardless of whether the BV or mouse reference genomes were used for the analysis (Fig. 5A), as well as by the comparison of gene expression profiles obtained with the two reference genomes (Fig. SD2C). Moreover, DEGs induced by PUUV and PHV showed a stronger correlation in tracheal cells (Fig. 5B), as shown by their distribution along the linear regression curve, than in renal cells (Fig. SD4) where several ISGs were induced at a higher level of FC by PHV than by PUUV (Fig. SD4B, right panel) as compared to a weak induction by PUUV and PHV in the tracheal cells (Fig. SD4A, right panel).

**Figure 5.**
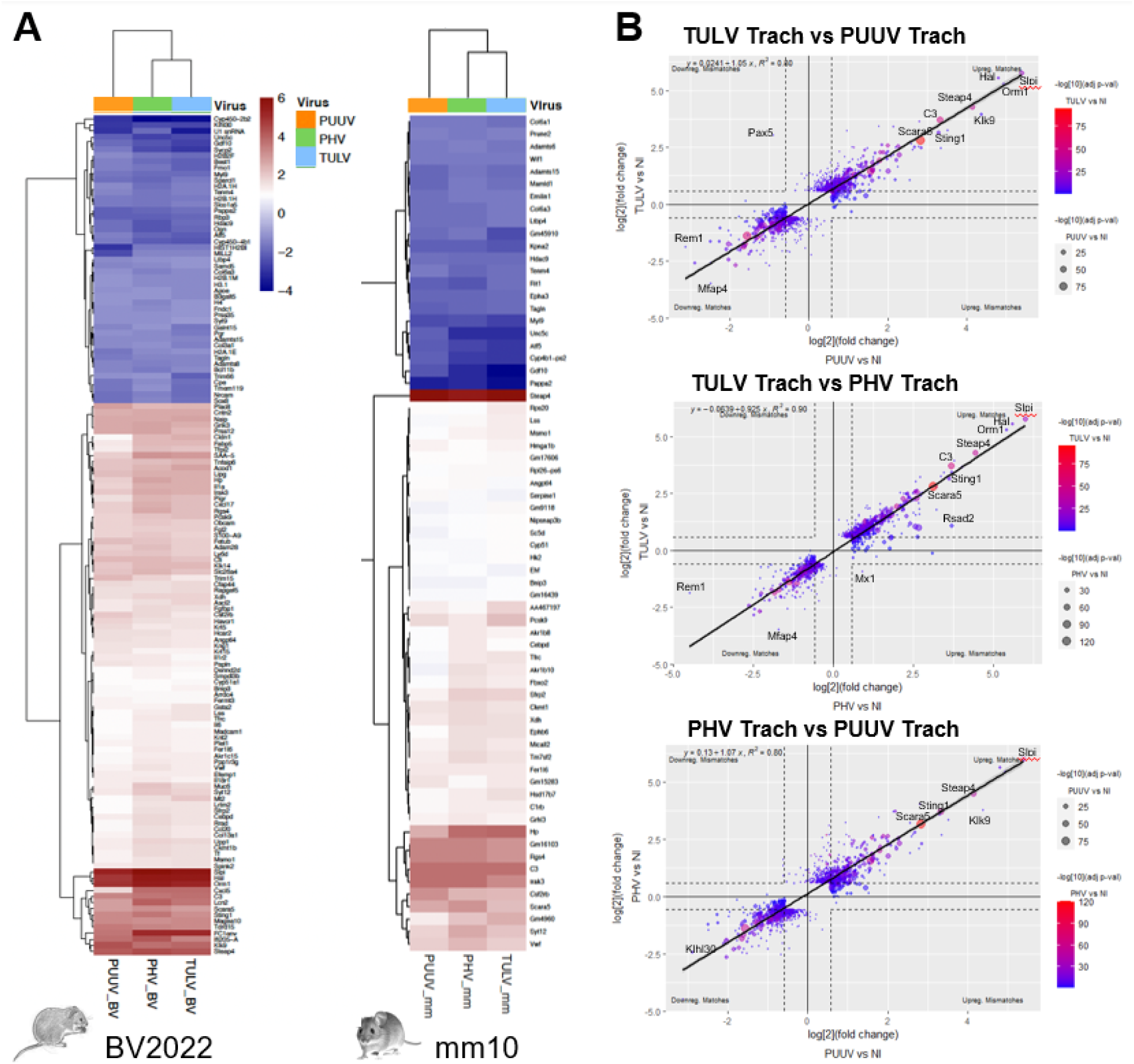
Gene regulation by orthohantaviruses infection of tracheal or kidney BV cells. Heatmaps (A) show the expression profiles of all differentially expressed genes (DEGs) in tracheal cells. Each column represents DEGs following infection with PUUV (dark orange), PHV (blue), or TULV (green) identified across two reference genomes, BV2022 and mm10 and each row corresponds to a gene name. Values are log2 fold changes for each gene relative to uninfected controls. Only genes present with all three viruses in tracheal cells are displayed. The color scale varies from dark red for highly upregulated genes to dark blue for highly downregulated genes, while white indicated no change. Genes and columns are clustered by expression similarity (Euclidean distance, complete linkage). Scatter plots (B) compared the log₂ fold change of host gene expression in tracheal cells (Trach) between TULV and PUUV (upper panel), TULV and PHV (middle panel) PHV and PUUV (lower panel) infections. For each panel, only genes displaying significant differential expression in at least one comparison (|fold change| ≥ 1.5 and adjusted p-value < 0.05) were retained. Each point represents a gene after filtering, and axes correspond to the log₂ fold change in the indicated infection conditions. Solid lines indicate zero fold change and dashed lines represent the fold-change threshold (|FC| = 1.5). Quadrants highlight genes showing concordant (matches) or discordant (mismatches) regulation between the two infections.

We therefore investigated whether certain signaling pathways specific to cells or orthohantaviruses could emerge from network analyses. DEGs from the innate immune response to infection and inflammatory processes were, as mentioned above, induced in different cell lines by PUUV and PHV orthohantaviruses (Fig. 4A). This was illustrated by STRING network representation. The antiviral immune response induced by PHV infection of renal, MyglaSWRecB, and respiratory cells, MyglaSWTrach and MyglaAEC#2, was predominant and showed strong network connectivity (Fig. SD5A) as compared to PUUV infection (Fig. SD5B). Besides, cell type-specific responses were observed as shown by the dot plot representations of biological processes (Fig. 6) by Gene Ontology enrichment. In addition to anti-viral defense, the DEGs that were activated by the three orthohantaviruses in infected tracheal cells included numerous membrane receptors, adhesion molecules and extracellular matrix (ECM) components with potential interacting functions, as well as transcripts involved in lipid metabolism and sterol pathways (Fig. 6A, Fig. SD6, Table SD1). STRING analyses also revealed virus-specific pathways in each cell type. For example, in MyglaSWTrach, PHV infection was associated with down-regulation of biological processes while ribosomal protein enrichment was observed after PUUV infection.

**Figure 6.**
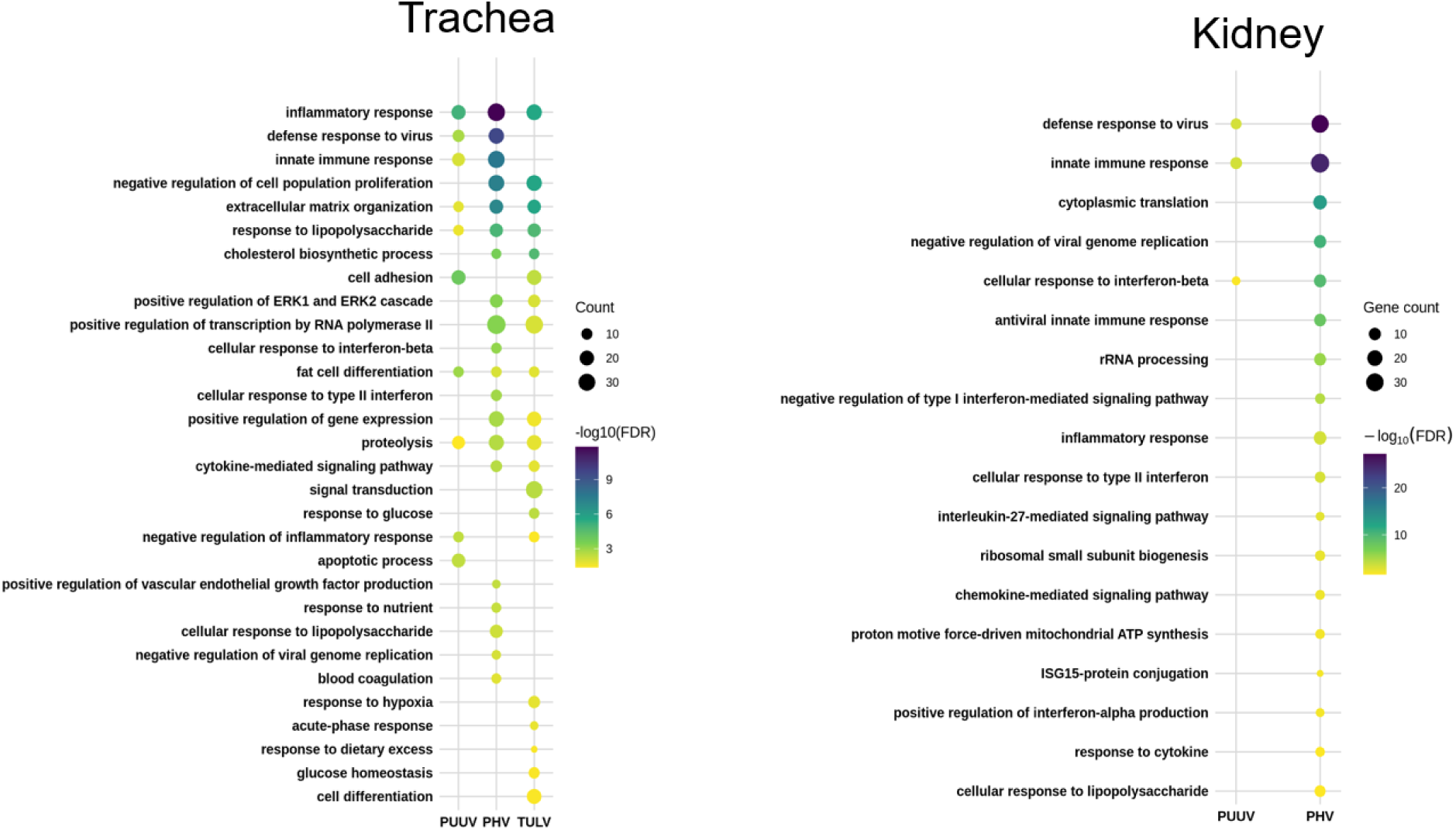
Comparative analyses of enriched pathways in tracheal and renal cells following orthohantavirus infection. Dot plots show significantly enriched biological process Gene Ontology terms identified using DAVID enrichment analysis. The left panels show analyses performed on differentially expressed genes from tracheal samples following infection with PUUV, PHV and TULV and the right panels kidney samples infected with PUUV and PHV. Only terms with FDR < 0.05 were retained. For each virus, a selection of the most significant enriched terms was ranked according to their FDR. Dot size represents the number of genes associated with each enriched term, while dot color indicates enrichment significance expressed as −log10(FDR). Terms are ordered according to the best FDR observed across all conditions.

The ECM and peripheral compartments, strongly represented in tracheal cells, could be targeted by orthohantaviruses for entry. In this context, several genes activated by the three viruses are of interest, such as an orthologue of FC1env, an endogenous retroviral membrane protein with fusion properties, Plac8 described as important for the entry of Sars-Cov2 into lung cells (43) and Hvacr1, a hepatitis C receptor (44). Integrins are described as receptors for orthohantavirus entry, but other membrane proteins could be involved (45–47). Although only moderately regulated, several mucosal and membrane proteins expressed in tracheal cells are worth mentioning, as they could participate in orthohantavirus entry or activate signaling receptors; for instance, proteins capable of binding to integrins, such as Madcam1 (α4β7) and Fermt3, upregulated, or Edil3 (α5β3) were downregulated by PUUV and PHV. Other adhesion molecules were activated by the three viruses such as contactin (Cntn2) and ephrin receptor B2 (Ephb2), by PHV and TULV such as Lrrc4, Vcam1, cadherin (Cdh16), or finally activated by PUUV and PHV (Timd4). A member of this family, Tim1, has been recently described as an entry receptor for HTNV (48). Molecules with lectin characteristics are important in host defense to microorganism and several lectins were differently expressed in MyglaSWTrach, such as Clec2e activated by the 3 viruses, Clec1a activated by PHV and TULV or Clec2d11 and Itln1 activated by PHV and PUUV, respectively. In infected kidney cells, Clec2h was activated by PHV and Ficolin1 by PUUV.

Slpi, a mucosal secreted inhibitor of protease, is of particular interest. It has multiple functions among which, it protects epithelial cells from microbial infection and could control pro-inflammatory cytokines (49, 50). This DEG induced by the three orthohantaviruses in tracheal cells, showed the highest score of fold change (FC > 40). Other membrane or secreted proteins like Steap4 (51), Orm1 or Hp having a role in acute phase reaction, or involved in inflammatory processes like CxCl5, Klk9 (52), Irak3 and Acod1 (53) were also strongly activated by the three orthohantaviruses. The case of the inflammatory process where some genes are activators, others are inhibitors and still others have a dual regulatory activity, illustrates the complexity of the interaction induced by orthohantaviruses.

Many studies have now documented the role of lipid in immune defense (54, 55). Many genes involved in lipid metabolism (Lipg, Pcsk9, Tm7sf2, Msmo1, Lss, Cyp51a1, Smpdl3b, Hsd11b1, Mtmr7, agpat4, Pla1a, Adh4, Sqle, Fdps, insig1, Scd4), lipid binding and transport (Fabp5, Sec14l4, Lbp, Abca17, Sec1413), or lipid clearance (Angptl4, Pla2r1), were induced mainly in tracheal cells by the three orthohantaviruses. The genes with highest FC score activated by the heterologous viruses were Pla2g2a, a phospholipase and Sncb, an inhibitor of phospholipase. Other genes of the lipid metabolism were downregulated in tracheal cells infected with PHV or TULV (Agmo, Cel, Tmem86a, Acot5, abcg4, Arsi, Mogat, Sult2b1, Hsd3b). A recent study (56) has described a downregulation of genes of the lipid metabolism in lung epithelial cells infected with the orthohantavirus Andes (ANDV). In kidney cells, only low-level activation of Cds1, a gene involved in lipid droplet growth was observed following PHV infection.

Interestingly, the three genes most strongly and exclusively activated by TULV in the tracheal cells were identified with the mouse database. They are involved in vesicle trafficking (Ap3s1), endosomal recycling and cytoskeletal organization (Wipf3). Moreover, podoplanin (Pdpn) by binding to a C-type lectin could modify cell morphology by inducing membrane protrusions (57, 58). This could explain the membrane structures previously observed in TULV-infected cells (14). Pdpn also interacts with the Mt1 metallopeptidase and the Clec2 lectin (59) which were also upregulated in MyglaSWTrach.

Eight subunits of H2, H3, H4 histones identified from Loc genes, were specifically downregulated in infected tracheal cells (Tab SD1A), as well as other histone modifiers. The deacetylase Hdac9, a histone modifier (60), could have a regulatory role in innate immunity and the control of viral infection. Furthermore, Bex2 known to regulate apoptosis and the cell cycle (61) was downregulated with the heterologous PHV and TULV in MyglaSWTrach cells. These different downregulations could enhance the antiviral response in tracheal cells by promoting apoptosis and/or by decreasing cell division and consequently viral replication.

Regarding the downregulation of DEGs, in MyglaSWRecB kidney cells, many different components of ribosomal subunits important for the regulation of mRNA translation were inhibited by PHV (Table SD2, Fig. 6 and SD5). Surprisingly, other ribosomal subunits components shared between three or two orthohantaviruses (Table SD1) were upregulated in infected tracheal cells, whereas only a few were specifically downregulated by each virus (Table SD3).

## Discussion

The increase in orthohantavirus outbreaks calls for a better understanding of the different outcomes of these viruses in their natural hosts. Their co-circulation may increase as a result of environmental damages and biodiversity loss, which can alter rodent habitats. This could favor emergence of new orthohantaviruses with increased virulence through reassortment of viral genomes, (2, 62, 63). Little is known about natural host-hantavirus interactions that underlie viral persistence, although rodents trigger an adaptive immune response to infection. This may reflect long-term co-evolution (3, 4). Understanding the host-cell restrictions that allow persistence of homologous orthohantaviruses, or elimination of heterologous orthohantaviruses at different stages of the viral cycle, including cellular entry, viral replication or particle budding is important for predicting their spread, particularly in the case of co-infection. We addressed this question by searching for factors potentially associated with such restrictions, comparing DEGs identified in different epithelial bank vole cell lines infected with the homologous orthohantavirus PUUV or the heterologous orthohantaviruses PHV and TULV.

PUUV is known to persistently infect bank voles. We have previously shown that infectious particles are released from the BV renal cell line, MyglaSWRecB, in which PUUV induces a low IFN response compared with that observed in human A549 cells. In contrast, PHV and TULV, which have evolved with different natural hosts, do not replicate efficiently and do not produce infectious particles in this BV cell line, whereas PHV induces a higher IFN response than PUUV. In that previous study, the *Mus musculus* reference genome was used to identify DEGs (13, 14). Here, we extended our analysis of gene regulation during orthohantavirus infection by including respiratory epithelial BV cells originating from the tracheal (MyglaSWTrach) and pulmonary alveolar compartment (MyglaAEC#2) and by using both, the mouse (mm10) and bank vole (*Myodes glareolus* NCBI 2022) databases for gene identification. Surprisingly, only 20% of the identified genes in tracheal infected cells and even less in renal cells (5% to 10%) were common to the two reference genomes (Fig. SD2B and SD3B). This limited overlap could be due to several factors (i) although well annotated, the mouse genome may not be sufficiently closely related to the bank vole genome and another well-annotated database, such as hamster or rat genome, might be more appropriate; (ii) although continually enriched with new genes, the updated BV genome is not yet complete.

The present analysis corroborates our previous data (14) and findings from the literature indicating that antiviral immune responses and inflammatory processes are also induced by hantaviruses in rodent reservoirs (12, 17, 64). However, we observed substantial variations in the antiviral response of infected BV cells, depending on the virus and cell type (Fig. 4). First, the weak response following PUUV infection in kidney cells, could reflect its persistence in bank voles, which may result either from strong viral antagonism of innate immunity or from a weak immune response that permits continued viral production. The low FC value of genes from the innate immune response in PUUV infected cells favor the latter possibility. Second, a different situation was observed with TULV, which is of low pathogenicity in humans, is endemic to the Old World and is associated with another vole species. Genes involved in the immune response were activated by TULV in tracheal cells, although no response was observed in renal cells. This could indicate that TULV, which is not efficiently produced in BV cells, is eliminated at an early stage of its viral cycle. Third, a strong IFN response in terms of number of genes, fold change magnitude, and network connectivity, was induced by PHV, which is non-pathogenic to humans, and endemic to vole’s species of the New World. This could correlate with the elimination of PHV by both, tracheal and renal BV cells through sequestration in autophagic vacuoles (14). Consistent with this interpretation, several genes involved in the regulation of apoptosis and autophagy, which could inhibit the budding and release of viral particles, were strongly activated in BV cells infected with PHV. Overall, these results support the notion that PHV, which has the potential to infect different cell types and species, induces a strong antiviral response and could be retained and degraded intracellularly in a host species with which it has not co-evolved. In contrast, TULV, which does not efficiently replicate in different cell lines (14), induces an antiviral response in tracheal cells but not in MyglaSWRecB or MyglaAEC#2 cells, suggesting that antiviral mechanisms may restrict TULV replication at an early stage, potentially during viral entry.

While it has long been thought that hantavirus infection can induce responses that are specific to both the virus and the cell type (11, 65–68), the underlying mechanisms have yet to be defined. Furthermore, it should be noted that most studies of the orthohantavirus-host interactions described in the literature are based on studies of human endothelial cells or monocytes, infected with virulent viruses that are highly pathogenic to humans, such as ANDV and HTNV (65, 67, 69, 70). The present study identified several major enriched pathways in bank vole cells that displayed clear cell-type-specific patterns. In MyglaSWRecB cells, in addition to the activation of immune responses and inflammation by PHV, several ribosomal proteins involved in RNA translation were downregulated. In MyglaSWTrach cells, different genes involved in antiviral defense including components of the IFN-independent STING pathway (33) were activated, as well as DEGs involved in ECM organization and lipid metabolism. In addition, downregulation of histones in these tracheal cells are consistent with recent studies highlighting the interplay of epigenetic reprogramming with antiviral immune responses (71, 72).

A recent transcriptomic analysis of the tropism of Old World (HTNV) and New World (ANDV, SNV) pathogenic orthohantaviruses (56) also highlighted regulation of genes shared by these three viruses, or specific to each of them. Several genes involved in innate immune response and inflammatory processes, described in that study were also identified in our study, while only one of the identified receptors, Havcr1, was found in both studies. The complexity of the virus-host interaction resulting from the great diversity of orthohantaviruses and host species may account for some discrepancies between studies. For instance, the cholesterol pathway, slightly upregulated in our study, was downregulated in the work of Jeyachandran et al. These differences may be related to stronger immune responses expected in human cells than in rodent cells. Genes belonging to the sterol regulatory pathway have also been described as required for ANDV entry in infected haploid human cells (73). Although, different genes may be identified, they remain of interest given the reported interplay between fatty acid and lipid metabolites and the innate immune response (27, 54, 55). As already mentioned, very few studies have explored orthohantavirus interactions with their natural hosts, largely because of the limited genomic information and the lack of experimental tools available for these species. Nevertheless, a transcriptomic analysis of splenic RNA from naturally infected long-tailed pygmy rice rats, the reservoir of ANDV, was performed to identify markers potentially associated with viral persistence (74). None of the regulated genes involved in immune responses or lipid metabolism, which have been identified using the *Mus musculus* reference genome, were shared with our infected bank vole cells, although some genes belonged to the same families. Another study has used lymph nodes from deer mice experimentally infected with ANDV which is associated with long-tailed pygmy rats and is cleared from deer mice. The authors concluded that a Th2 skewed-adaptative immune response played a predominant role in these infection outcomes (75). Altogether, these transcriptomic analyses highlight the complexity of pathway regulation in orthohantavirus-infected-bank vole cells and the difficulty of determining which cellular model is most appropriate for studying specific aspects of virus-host interactions.

It should also be noted that other types of RNA, including snRNA and rRNA as well as numerous long non-coding RNAs (lncRNA) were sequenced in the BV cells. Their functions are unknown and they were therefore removed from the gene lists presented in the tables. However, lncRNAs can act as regulatory factors and some are known to regulate inflammation (76, 77) or ISGs expression (78) as shown in HTNV-infected cells (79). Further analysis of these lncRNAs may therefore provide additional insights into regulation of the antiviral response in BV cells.

Our work has identified genes not yet described in the antiviral response induced in rodents infected with orthohantaviruses, including genes potentially involved at different stages of the viral cycle, which may contribute to the different outcomes observed following infection. The use of different cell types also provides information on cell type-specific responses that could influence viral spread within an organism. Tracheal epithelial cells, which are among the first cells encountered by aerosolized viruses, showed a broad transcriptional response involving genes associated with stress response, lipid metabolism, ECM remodeling and potential mucosal receptors. Among these genes, some are known to protect epithelial tissue, which represents a first barrier to viral infection and may limit viral dissemination to other organs. Finally, potential receptors have been identified in BV cells such as members of the BTN family (80) among which BTN3 has been suggested to have zoonotic relevance for influenza virus (81). Different types of membrane proteins that could serve as receptors or co-receptors for viral entry have been documented in small mammals and may contribute to inter-species transmission (82). The present transcriptomic analyses provide additional information on the great diversity of orthohantavirus interactions with their natural host, both for orthohantaviruses specifically associated with bank voles and for heterologous orthohantaviruses capable of infecting a “non-natural” host. Beyond their relevance for understanding the mechanisms underlying orthohantaviruses persistence and dissemination in host reservoirs, the identification of potential receptors and host restriction factors may help improve our understanding of the determinants of orthohantavirus spillover potential.

## Acknowledgements

We are grateful to Isabella Eckerle, Christian Drosten and Marcel Muller (Charité Berlin) for providing the different Mygla cell lines generated within the EVAg program.

## Author contributions

KB performed enrichment analyses and representation and contributed to the writing

TC performed the RNA-seq analysis and contributed to the writing.

ET performed technical procedures for RNA sequencing

MM validated RNA-seq analysis and contributed to the writing

JCM acquired funding and contributed to the writing

ME conceptualized and supervised the work and wrote the original manuscript

GG conceptualized the work, performed analyses and contributed to the writing

## Supplementary data

**Table SD1:** Genes regulated in MyglaSWTrach cells infected by the 3 orthohantaviruses (A) or common to 2 out of 3 orthohantaviruses (B).

**Table SD2:** Genes regulated in MyglaSWRecB cells, common to PUUV and PHV (A) and unique to PHV (B) or PUUV (C).

**Table SD3:** Genes regulated in MyglaSWTrach infected cells, unique to PHV (A), PUUV (B) or TULV (C).

**Figure SD1: Quality of the RNA-seq**. The histogram in (A) shows the number of reads that could be assigned to genes of the bank vole NCBI2022 database for the different samples of non-infected (NI) and PUUV, PHV or TULV infected cells as indicated. For each condition in each indicated cell lines, 4 samples were sequenced and bars correspond to the mean of the number of reads with the standard deviation represented by the error bars. The 3D projection of the PCA analysis of the different samples is shown for MyglaSWRec (B) and MyglaSWtrach (C). The histogram (D) shows the number of reads that could be assigned to genes of the *Mus musculus* mm10 database for MyglaSWTrach infected by the 3 orthohantaviruses and the clustering of the samples by the 3D projection of the PCA analysis (E)

**Figure SD2. Comparison of DEGs identified with bank vole or mouse databases, in MyglaSWTrach infected cells.** In (A), the Venn diagrams show the percentage of genes identified with the BV database (left panel) and the mouse database (middle panel) specific or shared by tracheal cells infected with PUUV, PHV and TULV. The heatmap of the right panel in (A) shows the expression profiles of all DEGs in tracheal cells infected with the three viruses across the two reference genomes (BV and mm). Rows represent DEGs, columns represent virus/genome combinations. Missing values were replaced by 0. Values are log2 fold changes scaled by row (scale = “row”). Columns are clustered using Euclidean distance and complete linkage. The annotation bar indicates the virus (PUUV, PHV, TULV) and the corresponding genome (BV, mm). The Venn diagrams in (B) compare the percentage of DEGs specific or common to the BV and mouse database identified in trach cells infected with PUUV (left panel), PHV (middle panel) or TULV (right panel).

**Figure SD3. Comparison of DEGs identified with bank vole or mouse databases, in MyglaSWRec infected cells.** The Venn diagrams in (A) show the percentage of genes identified with the BV database (left panel) and the mouse database (right panel) specific or shared by tracheal cells infected with PUUV or PHV. The Venn diagrams in (B) show the percentage of DEGs specific or common to the BV and mouse database identified in renal cells infected with PUUV (left panel) or PHV (right panel).

**Figure SD4. Comparison of host transcriptional responses to distinct orthohantavirus infections in tracheal and renal cells.** Comparison of host gene expression in PUUV vs PHV is shown in (A) for tracheal (Trach) cell infection and in (B) for kidney cell (Rec) infection. Scatter plots (left panels) of the log₂ fold change show only genes displaying significant differential expression in at least one comparison (|fold change| ≥ 1.5 and adjusted p-value < 0.05). Solid lines indicate zero fold change and dashed lines represent the fold-change threshold (|FC| = 1.5). Quadrants highlight genes showing concordant (matches) or discordant (mismatches) regulation. Point color represents the statistical significance of differential expression in one infection condition (−log₁₀ adjusted p-value), while point size reflects the significance in the other condition. A linear regression line illustrates the correlation between transcriptional responses. The plots (right panels), highlighting genes contributing to the antiviral response, compare the log2 FoldChange of different genes regulated by PUUV versus PHV infections in Trach and Rec cells.

**Figure SD5. Network analysis of enriched pathways in the different bank vole cells infected with orthohantaviruses.** Gene ontology of biological processes enriched pathways for genes of the immune system (red), ISG15 antiviral response (blue) and lipid pathway (green) were obtained using the STRING online server. Genes from other pathways are in white. In (A), network connectivity is shown for genes induced by PHV in tracheal cells (left panel), renal cells (middle panel) and lung alveolar cells (right panel), in (B) the connectivity for genes induced by PUUV in tracheal cells (left panel) and renal cells (right panel) and in (C) for genes induced by TULV in tracheal cells. DEGs with (|fold change| ≥ 4 and adjusted p-value < 0.05) were considered for the analysis.

**Figure SD6 Network analysis of enriched pathways shared by different orthohantaviruses in tracheal infected cells.** Network connectivity of DEGs in MyglaSWtrach either common to the infection with PUUV, PHV and TULV (A) or shared only following PHV and TULV infection (B) was obtained using STRING. Gene ontology of biological processes is shown in red for genes involved in host defense response, in green for negative regulation of biological process, in blue for functions involving extracellular region and in yellow for cholesterol biosynthetic processes. The analysis included DEGs with |fold change| ≥ 2 and adjusted p-value < 0.05.

## Data availability

The RNA-seq data set 2 and 3 generated as part of this work were deposited to ENA (accession E-MTAB-16656).

