## Supplementary material for "Comparative transcriptomic response of bank vole kidney and respiratory epithelial cells to distinct orthohantaviruses": Table 1

**Table 1** : Characteristics of the RNAseq used for gene identification

| A) | MyglaSWRecB vs BV2022^4^ | | | | MyglaSWTrach vs BV 2022^4^ | | | |
| --- | --- | --- | --- | --- | --- | --- | --- | --- |
|  | NI | PUUV | PHV | TULV | NI | PUUV | PHV | TULV |
| Total reads^1^ | 28.9M  (sd 1.9) | 30.2M  (sd 1.9) | 28.3M  (sd 3.8) | 26.7M  (sd 7.7) | 31.3M  (sd 2.6) | 32.7M  (sd 5.2) | 35.4M  (sd 4.8) | 33.9M  (sd 2.9) |
| Assigned reads^1^ | 22.5M (sd 2.0) | 24.1M  (sd 1.5) | 22.1M  (sd 2.4) | 22.1M  (sd 8.7) | 25.0M  (sd 2.2) | 26.0M  (sd 4.42) | 27.9M  (sd 3.8) | 26.9M  (sd 2.3) |
| all retained genes^2^ |  | 687  Up 433  Down 254 | 1756  Up 1096  Down 660 | 4  Up 1  Down 3 |  | 4292  Up 2098  Down 2194 | 5463  Up 2715  Down 2748 | 5513  Up 2707  Down 2806 |
| Significantly up-regulated^3^ |  | 48  73% | 183  72% | 1  25% |  | 189  62% | 383  70% | 307  60% |
| Significantly down-regulated^3^ |  | 18  27% | 71  28% | 3  75% |  | 119  38% | 162  30% | 202  40% |
| Significantly regulated |  | 66 | 254 | 4 |  | 308 | 545 | 509 |

| B) | MyglaAEC#2 vs BV 2022^4^ | | | | MyglaSWTrach vs mm10^4^ | | | |
| --- | --- | --- | --- | --- | --- | --- | --- | --- |
|  | NI | PUUV | PHV | TULV | NI | PUUV | PHV | TULV |
| Total reads^1^ | 39.1M  (sd 10) | 27.4M  (sd 8.7) | 22M  (sd 6.7) | 33M  (sd 9.4) | 31.3M  (sd 2.2) | 32.7M  (sd 4.5) | 35.4M  (sd 4.2) | 33.9M  (sd 2.5) |
| Assigned reads^1^ | 30.6M  (sd 7.8) | 21.6M  (sd 6.8) | 17.1M  (sd 5.7) | 25.2M  (sd 6.6) | 7.0M  (sd 0.5) | 7.2M  (sd 0.9) | 7.9M  (sd 0.9) | 7.5M  (sd 0.8) |
| all retained genes^2^ (p<0.05) |  | 2  Up 1  Down 1 | 12 Up 12  Down 0 | 0 |  | 1620  Up 814  Down 806 | 2323  Up 1141  Down 1294 | 2577  Up 1283  Down 1294 |
| Significantly up-regulated^3^ |  | 0 | 2 | 0 |  | 68  48% | 189  71% | 187  67% |
| Significantly down-regulated^3^ |  | 0 | 0 | 0 |  | 75  52% | 79  29% | 93  33% |
| Significantly regulated |  | 0 | 2 | 0 |  | 143 | 268 | 280 |

^1^-“M” correspond to millions for the number of reads ; ^2^-all genes with a pvalue < 0.05 were retained and ^3^- among them genes detected with a fold change (FC) >2 were considered as differentially expressed in infected cells compared to non-infected cells; ^4^- BV stand for the bank vole (*Myodes glareolus*) NCBI 2022 data bank and mm10 for the *Mus musculus* bank

| C) | Number of unique genes | | | Number of common genes | | | | Number of total genes | | |
| --- | --- | --- | --- | --- | --- | --- | --- | --- | --- | --- |
| Infected  cells | PUUV | PHV | TULV | PUUV PHV | PHV TULV | PUUV TULV | PUUV PHV TULV | PUUV | PHV | TULV |
| MyglaSWTrach FC 2 & >2 | 82  10.5% | 169  21.6% | 121  15.4% | 24  3.1% | 186  23.7% | 36  4.6% | 166  21.2% | 308 | 545 | 509 |
| MyglaSWRec FC 2 & >2 | 37  12.5% | 226  76.6% | 3  1% | 28  9.5% | 0  0% | 1  0.3% | 0  0% | 66 | 254 | 4 |
