## Supplementary material for "Comparative transcriptomic response of bank vole kidney and respiratory epithelial cells to distinct orthohantaviruses": Table 2

**Table 2**: Genes common to different cell lines and or viruses

|  | **Mygla** | | | | | |  | **Function** |
| --- | --- | --- | --- | --- | --- | --- | --- | --- |
| **Genes** ^1^ | **SWTrach** | | | **SWRecB** | | **AEC^3^** | **A549^4^** |  |
|  | **PUUV** | **PHV** | **TULV** | **PUUV** | **PHV** | **PHV** | **PUUV** |  |
| **Upregulated genes ^2^** | | | | | |  |  |  |
| *Ddx60* | 10.4 | 53.9 | 13.8 | 3.5 | 53.3 |  | 4.6 | RNA helicase: promoting IFN via RIG & MDA5 |
| *Rsad2* | 4.5 | 13 | 2.0 |  | 23.3 | 1.02 | 20.4 | IFI antiviral defense: inhibitor of viral replication (budding) |
| Lgals9 | 4.3 | 4.7 | 3.9 |  | 6.7 | 1.01 |  | Galectin, modulating cell-cell and cell-matrix interaction |
| *Ifit3* | 3 | 5.9 | 2.0 |  | 13.8 |  | 6 | IFI- antiviral defense Inhibitor of viral replication |
| Cxcl10 | 2.2 | 5.3 | 2.2 |  | 2.2 |  | 18.1 | IFγ induced -proinflammatory cytokine; role in viral infection (Immune cell migration) |
| *Pla2g5* | 4.3 | 29.2 | 19.6 |  | 7.8 |  |  | **PLA2, lipid hydrolysis/mb lipid remodeling (host defense)** |
| Zbp1 | 2.1 | 4.9 | 2.8 |  | 11.1 |  |  | II sensor/ mediate pyro, necro and apoptosis, viral defense |
| Cfb | 3.0 | 3.5 | 2.9 |  | 4.4 |  |  | Complement factor B from alternate complement pathway |
| *Ube2l6* | 2 | 3.1 | 2.7 |  | 3.9 |  |  | Ubiquitination of proteins such as ISG15, p53, proteasome degradation |
| *Ifit1* | 2.4 | 5.6 |  | 2 .1 | 24.3 | 1.58 | 12.1 | IFI Inhibition of viral replication and translation |
| *Cmpk2* | 2.0 | 6.1 |  |  | 6.5 |  | 15.1 | Mitochondrial kinase. Antiviral host restriction factor, acts independently of IFN, inhibition of viral replication |
| *Ifit2* | 2.4 | 6.4 |  |  | 13.6 |  | 4.2 | IFI-antiviral defense; apoptosis; Inhibition of viral transcription |
| Mnda |  | 6.3 | 4.1 | 2.4 | 41.8 |  |  | Myeloid nuclear diff antigen similar IFI211 and Ifi205 genes |
| Tgtp2 |  | 6 | 3.8 | 2.9 | 26.8 |  |  | GTPase: IFNγ induced; antiviral= Irgb6 |
| *Oasl* |  | 11.5 | 2.4 |  | 14.3 | 1.05 |  | Antiviral activity independent of RNAse L (viral replication) |
| *Pla2g4a* |  | 2.6 | 2.4 |  | 2.3 |  |  | **Cyt PLA2/hydrolysis of mb PL/regulates inflammation** |
| FLO11-L |  | 2.1 | 2.0 |  | 3.4 |  |  | Flocculation protein |
| *Mx1* |  | 2.6 |  | 2 | 13.7 | 1.9 | 24.9 | IFI antiviral response |
| *Iigp1* |  | 6.5 |  | 2.1 | 37.0 | 1.1 |  | IFI GTPase, resistance to intracellular pathogens |
| Rtp4 |  | 3.3 |  | 2.4 | 15.4 | 1.6 |  | Receptor transporter (traffic) /IFI suppressor of viral replication |
| Oas2 |  | 5.8 |  | 2.2 | 16.9 |  | 19.5 | IFI antiviral response: inhibition of viral replication |
| Mx2 |  | 2.1 |  | 2.1 | 13.2 |  | 15.7 | IFI, antiviral response |
| Ifi203 |  | 3.3 |  | 2.3 | 16.9 |  |  | IFI Activation of innate immune response |
| IFI44L |  | 3.6 |  |  | 15.0 | 1.8 | 140.8 | ISG, IFI Regulator of antiviral response |
| *Irgm1* |  | 4.9 |  |  | 57.7 | 1.7 |  | GTPase: IFNγ induced, pro inflammatory cytokines, autophagy |
| Gbp6 |  | 3.2 |  |  | 14.4 | 1.54 |  | GTPase, defense response to IFN type I and II, |
| *Usp18* |  | 3.5 |  |  | 11.9 | 1.4 |  | Negative regulator of IFN-I signaling pathway; ISGylaton |
| Bst2 |  | 6.3 |  |  | 9.4 | 2.1 |  | IFI antiviral restriction factor, tetherin, blocks mb release |
| *Oas1a* |  | 11 |  |  | 7.4 | 2.8 |  | IFI Defense response to viruses, apoptosis,… |
| *Irf7* |  | 2.0 |  |  | 6.5 | 1.3 |  | IFN regulatory transcription factor |
| GBP4 |  | 3.6 |  |  | 10.3 |  | 26.7 | IFI GTPase; cellular response to IFNγ/regulate-antiviral response |
| Ccl5 |  | 4.5 |  |  | 2.1 |  | 4.7 | Chemokine; immunoregulatory and inflammatory processes, processed forms =Rantes |
| Samd9 |  | 2.6 |  |  | 4.3 |  | 4.2 | TNFα signaling, endosome fusion/cell proliferation and apoptosis |
| Samd9l |  | 4.1 |  |  | 3.1 |  | 4.3 | Innate immune defense to viral infection; endosome fusion |
| Ccdc194 |  | 9.3 |  |  | 12.4 |  |  | Coiled coiled domain containing protein; no info |
| Oasl2 |  | 3.4 |  |  | 11.1 |  |  | IFI Antiviral activity dependent of RNAse L |
| Apobec1 |  | 8.5 |  |  | 5.9 |  |  | Deaminase, mRNA editing enzyme; epigenetic regulator |
| Isg15 |  | 3.0 |  |  | 9.1 |  |  | Ubiquitin like modifier; IFNα/β induced, chemotactic to neutrophils, antiviral activity. |
| *ly6a* |  | 3 |  |  | 5.7 |  |  | Lymphocyte antigen; T cell activation |
| C2 |  | 2.2 |  |  | 8.8 |  |  | Serum protein part of the complement pathway |
| GBP5 |  | 4.3 |  |  | 6.9 |  |  | GTPase /Nlrp3 activator /II & inflammation/antiviral |
| Ifi27l2a |  | 3.9 |  |  | 5.0 |  |  | IFI, negative regulation of NR4A 1-3, IFNa inducible |
| Nlrc5 |  | 2.2 |  |  | 4.6 |  |  | NfKb, IFN-I and II regulation, control antiviral defense |
| *Mpeg1* |  | 2.8 |  |  | 2.3 |  |  | Macrophage expressed (perforin), IIR |
| *Dtx3l* |  | 3.6 |  |  | 3.3 |  |  | Ubiquitin ligase; IFI response to virus via PARP9 |
| *Irf9* |  | 2.2 |  |  | 3.5 |  |  | IFN regulatory transcription factor |
| *Ly6e* |  | 2.1 |  |  | 3.4 |  |  | Ly antigen, regulation of T cell, TCR, restriction of virus entry |
| Apol3 |  | 2.1 |  |  | 2.9 |  |  | **Apolipoprotein, Lipid movement & binding; TNF inducible** |
| Inka1 |  | 2.4 |  |  | 2.6 |  |  | Inhibitor of ser/thr protein kinase PAK4, Inka box actin regulator |
| Calhm6 |  | 2.1 |  |  | 2.3 |  |  | IRF3-Dependent NK-Activating Molecule; pathogen and pro-inflammation induced its relocation |
| Lrrc25 |  | 2.7 |  | 2.2 |  |  |  | Inhibits RLR mediated IFN by RIGI degradation (autophagy) |
| Ulbp1 |  |  | 2.5 |  | 2.0 |  |  | Unusual MHC-I ; mediates NK cell cytotoxicity |
| Cxcl11 |  |  |  |  | 5.5 |  | 22.5 | Chemokine IFIγ induced; chemotactic for Il activated T cells, PMN; binds Cxcr3 |
| *Ifih1* |  |  |  |  | 4.3 |  | 5.9 | MDA5, RLR signaling |
| *Ddx58* |  |  |  |  | 4.3 |  | 4.1 | RNA helicase promoting IFN signaling |
| *Epsti1* |  |  |  |  | 3.8 |  | 4.4 | Epithelial stromal interaction/MΦ activation |
| Xaf1 |  |  |  |  | 2 |  | 243 | IFI - regulator of IAP (apoptosis inhibitor), mediatesTNFα apoptosis |
| **Downregulated genes ^2^** | | | | | |  |  |  |
| Hist1h2ah |  | -2.1 | -2.0 |  | +2.2 |  |  | Histone H2A type 1H (H2A1H) |
| *Ackr4* | -3.8 | -3.1 | -4.0 |  |  |  | +5.8 | Atypical chemokine receptor |
| *Acta2* | -3.0 | -3.2 | -3.4 |  | +2.0 |  |  | Actin protein/intercellular signaling/cell motility |
| U1 SncRNA | -8.0 | -3.7 | -11.0 | -30.4 |  |  |  | U1 small nuclear RNA spliceosome |
| Cfap61 |  | -3.1 |  | -2.0 |  |  |  | Cilia and flagella assembly |
| *Rps17* |  |  | -7.9 |  | -2.3 |  |  | Small ribosomal subunit; protein synthesis |

^1^ Significant genes (p-adjust <0.05) with at least one condition exhibiting a fold change FC> or = 4 are shown. When identified only in mouse genome dataset they appear in red and when identified in both mouse and bank vole databases, they are in italic. ^2^ FC values correspond to the DEGs in infected cells versus non infected cells using analysis referring to BV genome. When higher values were observed in *Mus musculus* analysis they are reported in green and in red when only found with the mouse genome. ^3^ although the FC of DEGs in MyglaAEC#2 cells are very low, they are significant and are therefore reported. ^4^ the FC value of DEGs in A549 human cells infected with PUUV are from a previous identification on human genome data and are added for information (Gallo et al. 2020).
